# Extended genomic regions flanking ultraconserved elements allow efficient species identification and intraspecific diversity assessment in coral

**DOI:** 10.64898/2026.08.25.743606

**Authors:** Alejandro Mateos, Peter F. Cowman, Tom Bridge, Yun Kit Yeoh, David Bourne, Yui Sato

**Affiliations:** College of Science and Engineering, James Cook University, Townsville, QLD, Australia; AIMS@JCU, James Cook University, Townsville, QLD, Australia; Natural Sciences, Queensland Museum Tropics, Townsville, QLD, Australia; Centre for Tropical Bioinformatics and Molecular Biology, Townsville, QLD, Australia; Australian Institute of Marine Science, Cape Cleveland, QLD, Australia

**Author notes:** Correspondence: Y. Sato.

**Keywords:** Population genomics, genomic species identification, genetic diversity, targeted capture

## Abstract

Genetically informed conservation is critically important to ensure that interventions benefit the population of interest. In corals, preserving genetic diversity and accurate species identification are crucial for the sexual propagation. While various methods exist for species identification and measuring intraspecific variation, obtaining and analysing molecular data that enables rapid yet informed decisions on broodstock choice and progeny quality assurance remains challenging. Here we present a novel approach towards resource effective intraspecific genetic profiling by targeting extended genomic regions around ultra-conserved elements (UCEs). By sorting loci by parsimony informativeness and using a locus window size as small as 5000 bp upstream and downstream of the UCE, we identified a subset of 500 UCE-associated loci that can accurately resolve phylogenetic relationships among species and assess intraspecific variation with accuracy comparable to a whole-genome dataset, while. This method was validated using existing population genomic data from six species of staghorn coral (*Acropora hyacinthus, Acropora tersa*, *Acropora pectinata*, *Acropora* sp. ‘VI-3’, *Acropora kenti* and *Acropora* cf. *spathulata)*. The phylogeny produced by the UCE subset is congruent with the phylogeny based on complete data. With the moderate number and length of target genomic region sizes providing a balance between resolution and sequencing effort, this study provides a proof-of-concept approach towards developing fast, scalable, and cost-effective workflows using a real-time long-read sequencers such as Oxford Nanopore Technologies. The methodology has the broad potential to be applied to support genetic assessment across taxa where taxonomic uncertainty is common, improving confidence in experimental frameworks and conservation decisions.

## Introduction

Restoration activities are expanding to address the global decline of reef ecosystems and enhance resilience by replenishing coral populations on impacted areas (Bayraktarov et al., 2019; Peixoto et al., 2024). Coral propagation approaches such as coral transplantation, gardening (transplantation after nursery phase), and micro-fragmentation, involve propagation of asexually-reproduced, clonal lineages, which results in low genetic diversity within replenished populations (Baums, 2008). Consequently, there is an increasing focus on developing sexual propagation methods where gametes derived from multiple spawning parents are fertilised to generate offspring coral juveniles that are released at a given restoration site (Boström-Einarsson et al., 2020). Sexual propagation produces genetic variation by crossing multiple parent genotypes while the offspring genetic diversity is limited by the number and diversity of parents (Chamberland et al., 2017; Dallmeyer-Drennen et al., 2026). Sexual propagation can therefore be suitable to conserve the genetic variation in propagated corals allowing adaptive potential to environmental change that may enhance population viability, thereby better preserving ecosystem stability and functions (Baums et al., 2019; Hein et al., 2021). The indicators evaluating the success of restoration programs are often focused on the number of coral fragments created and post-transplantation growth and survival rates; however, other metrics such as genetic diversity are often overlooked (Hein et al., 2020). The application of genomic tools into restoration initiatives has been recognized as one of the critical components to address key knowledge gaps regarding the effectiveness of restoration activities (Banaszak et al., 2023; Baums, 2008). However, the lack of well-developed tools and workflows applicable to coral restoration presents one of the challenges to gain knowledge regarding the genetic and genomic landscape of populations and species subjected to restoration programs (Boström-Einarsson et al., 2020; Hein et al., 2021; Hughes et al., 2023).

Correct taxonomic identification of broodstock corals is key for captive coral propagation programs (Baums et al., 2019). For example, mistakenly breeding different species can reduce production of larvae due to hybrid incompatibility (Furukawa et al., 2024; Ramírez-Portilla et al., 2022; Willis et al., 1997). Coral identification is difficult however because traditional taxonomic resources (e.g. Veron, 2000) are based entirely on macromorphological characters that molecular phylogenetic data have shown do not reflect species boundaries or systematic relationships (Bridge et al., 2024; Connelly et al., 2026; Kitahara et al., 2016; Oury et al., 2023; Ramírez-Portilla et al., 2022; Rassmussen et al., 2025). Genomic data are effective to address issues associated with misidentified, pseudo-cryptic and cryptic species (Riginos et al., 2024).

In recent years, numerous approaches have been developed for resolving phylogenetic relationships between coral species and characterising within-species (intraspecific) genetic variation. Sequencing of ultraconserved elements (UCEs) - short, genome-wide, highly conserved regions among wide taxa (100∼200 bp-long ‘core’ regions; (Bejerano et al., 2004; Cummins et al., 2024) - and their flanking regions containing SNPs is useful for the reconstruction of phylogenies for resolving species at various evolutionary timeframes, with high cross-species applicability (Faircloth et al., 2012; Duckett et al., 2023). Using target enrichment of UCE loci for species boundary delimitation offers approaches to reduce analysis time for rapid species identification with accuracy (Erickson et al., 2021) and it has been applied to corals. For example, a bait set (DNA sequences designed to capture and isolate target genomic regions) targeting UCEs for Anthozoan phylogenomics was developed by Quattrini et al. (2018). Recently, this bait set was refined to specifically target UCEs within hexacorals with further specificity to Scleractinia (Cowman et al., 2020), and octocorals (Erickson et al., (2021). The development of these probe sets resulted in the revision of taxonomy at the species level for several complex coral taxa (Bridge et al., 2024; Connelly et al., 2026; Oury et al., 2025; Ramírez-Portilla et al., 2022; Rassmussen et al., 2025).

Within-species genotypic information of corals in breeding programs is also effective to avoid clonal genotypes used in the broodstock, causing skewness in parental genotype presentation in fertilisation, and to ensure high genetic diversity of broodstock corals and progenies produce. For intraspecific genotyping, current mainstream methodologies include the use of targeted sequencing to identify genome wide SNPs in genomic markers (e.g., microarrays and restriction-site associate DNA sequencing – RAD-seq), allowing assessment of genotypic variation at a large number of loci across the genome cost-effectively (Grover & Sharma, 2016). Approaches to assess intraspecific genetic variation are well developed and widely applied in population genetics and genomic studies of many coral taxa, using microsatellite genotyping (Van Der Ven, Flot, et al., 2021; Van Der Ven, Heynderickx, et al., 2021; Zayasu & Suzuki, 2019) and RAD-seq (Aurelle et al., 2021; Combosch & Vollmer, 2015; Iguchi et al., 2019; Torquato et al., 2022), while they do not usually include the targeted capture of UCEs. Data from these approaches are typically “single-use”, as microsatellites are taxon-specific, and the challenge of locus drop out in RAD-seq datasets means that they cannot be combined across species or studies (Bonito et al., 2021). The versatility of assays across taxa may be expected by anchoring SNP-based genotyping to genomic regions associated with loci that are found across the tree of life, including UCE regions.

In coral propagation programs, the collection of corals for spawning events and larvae development all happen inside small time windows (days to weeks) (Shlesinger & Loya, 2019), making rapid identification of broodstock often unachievable by conventional sequencing technologies generally provided by outsourced services. These challenges provide an opportunity to leverage recent advances in portable real-time sequencing with targeted long-read sequencing such as the Oxford Nanopore Technologies platform (Wang et al., 2021). Starting from DNA extraction, this platform allows for real time sequencing and can take less than 72 hours to obtain long (>10 kbp) contiguous reads that simplify assembly, increase mapping certainty, and improve variant detection (Amarasinghe et al., 2020; Kolmogorov et al., 2023). These characteristics warrant testing of a concept of a long-read real time sequencing approach that is suitable for use in the coral aquaculture timeframe. The first step in the development of such a rapid coral genotyping tool is to characterise efficient genomic targets that robustly identify genetic variation for species-level identification and intraspecific resolution for broodstock selection. This study tests the feasibility of combining the species-level phylogenetic resolution provided by UCE-derived phylogenomics and the capture of intraspecific variation with SNP loci in extended flanking regions around UCE loci (Fig. 1), by comparing with a whole genome sequencing data set within the ecologically dominant but taxonomically challenging genus *Acropora*. Furthermore, for the possibility of increasing data- and cost-efficiency, we test whether a subset of UCE-associated loci can be focused to verify species identity and to test for patterns in intraspecific genetic variation.

**Figure 1.**
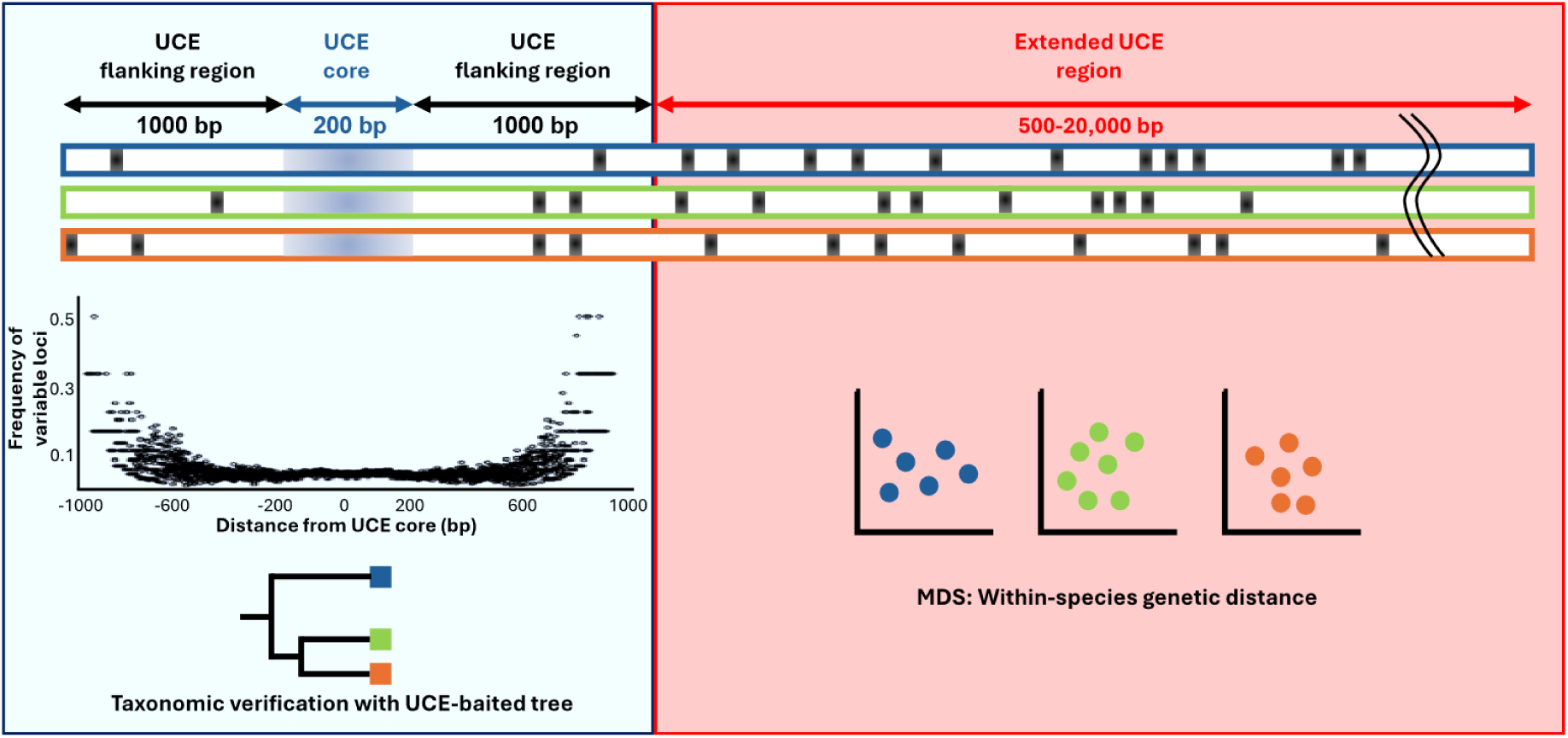
Schematic illustration of the study concept. UCE loci (∼200 bp core ± ∼1kbp flanking regions) are genomic regions that are used to capture phylogenetic information to resolve taxonomic identity. Each UCE loci was extended further upstream and downstream by ± 500 bp up to ± 20kbp (total locus window size of 1 kbp to 40 kbp) using each UCE core as an anchor to generate “extended UCE regions”, from which SNPs were extracted to examine the capacity to resolve intraspecific variation. The blue-, green-, and orange-coloured bars each represent an individual, the black ticks along each bar represent SNP loci distributed across the extended UCE region. The plot on the left panel represents the frequency of variable loci along the UCE locus in multi-species context, with variation increasing as it moves either side of the core.

## Materials and Methods

### Data source

Low-coverage whole genome sequencing (lcWGS) data derived from paired-end Illumina short reads were initially screened from existing public datasets, including 639 individuals from four tabular Acropora species (identified as *A. hyacinthus, A. tersa*, *A. pectinata*, and *A*. sp. ‘VI-3’ currently an undescribed species by Rassmussen et al. 2025; data source – Naugle et al., 2024; approximately 5-10x coverage), 698 *A. kenti* (Matias et al., (2023); BioProject PRJNA849642; approximately 5x coverage), and 831 *A.* cf. *spathulata* individuals (Denis et al., 2026; approximately 10x coverage). The data of *A. hyacinthus, A. tersa*, *A. pectinata*, *A*. sp. ‘VI-3’, and *A.* cf*. spathulata* were derived from individuals from 12 locations across the Great Barrier Reef (GBR) and the *A. kenti* data are from 26 locations across the GBR in the original studies. Subsequently, individual genome data were selected based on at least 4.0x coverage to ensure sensitivity and accuracy in variant detection and read mapping. In total, this study assessed 100 samples each for the *A.* cf*. spathulata* and *A. kenti*, while a smaller number of samples were studied for *A. tersa* (n=60), *A. pectinata* (n=20), *A*. sp. ‘VI-3’ (n=18), and *A. hyacinthus* (n=20), based on data availability.

### Read quality control and extraction of UCE loci for phylogenetic verification

A subset of samples (five for *A. tersa, A. pectinata, A. hyacinthus*, two for *A.* sp. ‘VI-3’, nine for *A. kenti*, and 14 for *A.* cf*. spathulata*) were selected for taxonomic verification based on read quality and largest file size. The reads were processed using fastp v.0.23.2 (Chen et al., 2018) with default values for adapter trimming, quality filtering, and base correction, and assembled using SPAdes Genome Assembler v.4.1.0 (Bankevich et al., 2012) with default parameters. The assembled contigs were then processed using a modification of the Phyluce v.1.7.3 pipeline (Faircloth, 2016), where we performed loci alignment outside of Phyluce v.1.7.3 with MAFFT v.7.0. The hexa-v2 UCE and exon bait set targeting 2,476 loci (Cowman et al., 2020) was matched to the *de-novo* assembled contigs at 70% identity and 70% coverage using *phyluce_asembly_match_contigs_to_probes*. The matching loci were then extracted using *phyluce_assembly_get_match_counts* and *phyluce_assembly_get_fastas_from_match_counts*. Loci from representative samples and type specimens (topotypes, collected from the from the same locality as the holotype, and paratypes, specimens not designated as holotypes but collected at the same location and time; housed in the collections of the Queensland Museum Tropics as part of the CoralBank database; provided Cowman & Bridge; https://www.museum.qld.gov.au/collections-and-research/projects/project-dig/research-projects/coral-bank) were added to the extracted UCE loci prior to alignment to aid in taxonomic verification. Singleton loci containing only one sample (i.e. non-informative loci) were filtered out before alignment. The aligned loci were then merged to the alignments from the Cowman et al. (2020) study (151 samples from the genus *Acropora*, one sample of *Isopora* cf*. brueggemanni* as an outgroup) with MAFFT v.7.0 (Katoh, 2002) using the *--add* and *--reorder* flags to align the new sequences to the existing alignments. The resulting concatenated alignments were edge-trimmed using *phyluce_align_seqcap_align*. Finally, alignment data matrices were generated using *phyluce_align_get_only_loci_with_min_taxa*, in which loci with more than 75% species occupancy were retained for downstream analysis.

For taxonomic verification, maximum-likelihood (ML) inference was run on the alignment using IQ-TREE v.2.3.6 (Nguyen et al., 2015). The analysis was performed with 1,000 ultrafast bootstrap replicates and a single best ML tree search on the concatenated alignments, all under the GTRGAMMA model (general time reversible gamma rate heterogeneity).

### Assessment of intraspecific genetic variation

Publicly available annotated scaffolds from whole genome resources published as ‘*Acropora hyacinthus’* (BioProject PRJEB81534; probable identity is *A. tersa;* pers. comm. T. Bridge, 2025), ‘*A. tenuis’* (Liew et al., 2016; probable identity is *A. kenti;* pers. comm. T. Bridge, 2025), and *A.* cf*. spathulata* (BioProject: PRJNA1013826) were used to generate sequence references encompassing loci of the core UCEs and their extended flanking regions for intraspecific variant identification (Fig. 1). These extended UCE region references were generated by capturing flanking regions both upstream and downstream of the UCE cores from each reference genome at six different lengths (± 500 bp, ± 1 kbp, ± 2 kbp, ± 5 kbp, ± 10 kbp and ± 20 kbp) to examine for the most resource-efficient sequence sizes. The selected extended region size range was guided by the standard size of a UCE-associated analyses (Faircloth et al., 2012; a core of approximately 500 bp and 500 bp flanking regions each side; a total of an approximately 1.5 kbp window size), and practical read lengths achieved by Oxford Nanopore sequencing (Amarasinghe et al., 2020; herein testing 2.5 kbp, 10.5 kbp, 20.5 kbp, 40.5 kbp). Specifically, the references were first generated using the Phyluce v.1.7.3 *harvest from genomes* pipeline, by matching the genome assemblies with the hexa-v2 bait set (Cowman et al., 2020) with *phyluce_probe_run_multiple_lastzs_sqlite*, and extracted with *phyluce_probe_slice_sequence_from_genomes* using the *--flank* flag to extend the lengths of each UCE’s flanking region (i.e., 500 bp ∼ 20 kbp either side of the UCE core).

SNPs in the extended UCE regions and genome wide SNPs were identified among all individuals using a lcWGS workflow (Modified from Sato et al., 2023 and Therkildsen & Palumbi, 2017). Modifications were made to accommodate the separation of algal symbiont DNA and to avoid false-positive SNP sites. The lcWGS sequences were prepared for mapping by deduplicating with fastuniq v.1.1 (Xu et al., 2012) to remove duplicate reads and reduce allele frequency biases from PCR artifacts, followed by adapter clipping with Trimmomatic v.0.33 (Bolger et al., 2014) to remove adapter sequences remaining from Illumina sequencing. The cleaned reads were then matched to algal symbiont genomes (Clade A: *Symbiodinium microadriaticum*, Aranda et al., 2016, PRJNA292355; Clade B: *Breviolum minitum*, Shoguchi et al., 2013, PRJDB732; Clade C: *S. goreaui*, Liew et al., 2016; Clade D: *Durusdinium sp.*, Shoguchi et al., 2021) and separated using bbsplit v.35.85 to discard sequences matching to algal symbionts followed by the removal of low-quality bases and overly short reads using Trimmomatic v.0.33 (Bolger et al., 2014) and merging of overlapping reads with FLASH v.1.2.11 (Magoč & Salzberg, 2011). The input reads were size capped to 1.8 Gb per sample to ensure similar mapping depths and improve SNP detection downstream. The processed lcWGS sequences were then mapped to the extended UCE region references, as well as the whole genome (WG) references for benchmarking. The sequences were then aligned to the references using bowtie2 v.2.5.4 (Langmead et al., 2009) and realigned with GATK v.3.8 (McKenna et al., 2010) for better SNP detection. This resulted in an average sample depth of approximately 3.0x for *A. hyacinthus, A. tersa*, *A. pectinata*, *A*. sp. ‘VI-3’ and *A.* cf. *spathulata;* and 1.0x for *A. kenti*. Mapping depth identification with ANGSD v.0.938 (Korneliussen et al., 2014) was performed across reference nucleotide sites to account for potential SNPs misidentified from erroneous mapping, owing to insufficient reads or cross mapping from repeat regions, thereby reducing the noise in SNP data. Upon assessment of global mapping depth distribution, SNP sites with minimum and maximum global read depths of 250-450 were searched for *A. hyacinthus, A. tersa*, *A. pectinata*, *A*. sp. ‘VI-3’, 33-160 for *A. kenti*, and 260-410 for *A.* cf*. spathulata*, respectively. Sites with >10 reads mapped in an individual sample were also excluded from the SNP search to exclude repeated regions while allowing high read depths from stochastic processes in the sequencing. SNPs were called by computing posterior genotype probabilities in a Bayesian approach with ANGSD v.0.928 (Korneliussen et al., 2014), assuming Hardy-Weinberg equilibrium (HWE) applied to sample allele frequency as a prior, with a SNP p-value<0.001, and a minimal minor allele frequency of 0.01. Sites extremely departed from HWE were excluded (p<0.001).

To characterize intraspecific genetic variation captured across the extended UCE loci and WG references, nucleotide diversity (*π*) (Nei & Li, 1979) and Watterson’s theta (*θ*) (Fu & Li, 1993) were estimated. Watterson’s estimator quantifies the number of polymorphic sites, while nucleotide diversity describes the average pairwise nucleotide difference between all sequences. The *π* and *θ* were estimated from the posterior genotype probabilities, based on per-site allele frequency likelihoods and the folded site frequency spectrum (SFS) using the *realSFS* command. All parameters were calculated separately for each of the six extended UCE region references as well as the WG reference.

To assess the resolution of genetic divergence among intraspecific individuals, pairwise genetic distances were estimated across the approaches, using ngsDist v.1.0.1 (Vieira et al., 2016) based on the ANGSD variant calling. Genetic distance matrices were then used for visualization with multidimensional scaling (MDS) using ggplot2 v.3.5.1 in R v.2024.12.1 (R Core Team, 2021). To evaluate among-species clustering, MDS was first performed for *A. tersa, A. pectinata, A*. sp. ‘VI-3’, *A. hyacinthus* together. To ensure within-species evaluation, this first MDS analysis was also used to identify the main clusters in *A. kenti* and *A*. cf. *spathulata*, and subsequent analyses were limited to one of the identified clusters.

The resolution of intraspecific divergence patterns was tested with Mantel tests, by comparing the genetic distance matrices derived from SNPs called from the extended UCE regions with those derived from the WG references. The Mantel tests were performed with 999 permutations using Pearson’s correlation coefficient (standard linear correlation) to calculate Mantel’s r, using the *mantel* function of the R package vegan v.2.6.4 (Dixon, 2003) and plotted using ggplot2 v.3.5.1.

### Loci subsetting and reduced loci analyses

As the number of targeted loci increases, so does the requirement for additional probes and amount of data needed to achieve sufficient sequencing depth per locus. To assess the effect of the number of UCE-associated loci on the species identification and intraspecific variation assessments, the above processes were repeated using the same parameters but based on subsets of UCE-associated loci. To enable the selection of UCE-associated loci focussing on the taxonomic resolution, a data frame was generated to select loci based on phylogenetic informativeness. The number of parsimony informative sites (i.e. the number of sites that have information that can reconstruct the evolutionary history of a group) per locus were obtained based on the ML tree containing all reference species and all samples tested above, using the *phyluce_align_get_informative_sites* phyluce v.1.7.3 program. The UCE loci were sorted by the descending number of informative sites to identify the top 100, 200 and 500 loci among those recovered consistently across the *A. kenti*, *A.* cf*. spathulata* and *A. tersa* references studied.

## Results

### UCE phylogenomics and taxonomic verification

The UCE search recovered a total of 1,781 non-overlapping UCE loci from the assembled contigs from *A*. *tersa*, *A. hyacinthus*, *A.* sp. ‘VI-3’, *A. pectinata*, *A. kenti*, and *A.* cf*. spathulata*. The mean number of loci per sample for *A*. *tersa*, *A. hyacinthus*, *A.* sp. ‘VI-3’ and *A. pectinata* was 1,570 ± 58 (mean ± SD; n = 17) and 1,506 ± 39 for *A.* cf*. spathulata* (n = 14). A lower number of loci were recovered for *A. kenti* (728 ± 227; n = 9).

The ML phylogenetic tree based on the above UCE loci were supported by the bootstrap values of 100 at 61.2% of the internal nodes (70.5% of nodes resolved with bootstrap ≥ 95), showing the six-clade topology of the *Acropora* genus consistent to Cowman et al. (2020); (100 bootstrap value for each of the six clades) (Supp. Fig.1). Each species tested in this study was reconstructed as reciprocally monophyletic (*A. tersa* n = 5, *A. hyacinthus* n = 5, *A.* sp. ‘VI-3’ n = 2, *A. pectinata* n = 5, *A. kenti* n = 9 and *A.* cf*. spathulata* n = 14; with bootstrap value ≥ 95). Additionally, the sample sequences derived from lcWGS were placed along the topotypes, paratypes or the representative control samples’ UCE loci sequences from CoralBank database.

### Intraspecific variation

MDS plots were used to visualize genetic divergence among conspecific samples based on genetic distances (Fig. 2). Overall clustering patterns were consistent across all extended UCE regions compared to the WG dataset in all species, with tighter grouping observed with increasing region size. MDS on *A*. *tersa*, *A. hyacinthus*, *A.* sp. ‘VI-3’ and *A. pectinata* showed consistent clustering across all extended UCE regions and was able to group the samples according to the four species (Fig. 2a), with the first two principal components respectively explaining ∼11% and ∼5% of the variance. For *A. kenti*, the first two components explained ∼16% and ∼2% of the total variance, respectively, across all extended UCE region sizes. The samples formed two clusters (Fig. 2b). Similar patterns were observed for *A.* cf*. spathulata*, with low variance explained by the first two dimensions (∼3% and ∼1.5%), while two clusters that, according to the metadata associated with these samples, reflect the genetic differences between individuals from the Southern GBR and the Central and Northern GBR (Fig. 2c).

**Figure 2.**
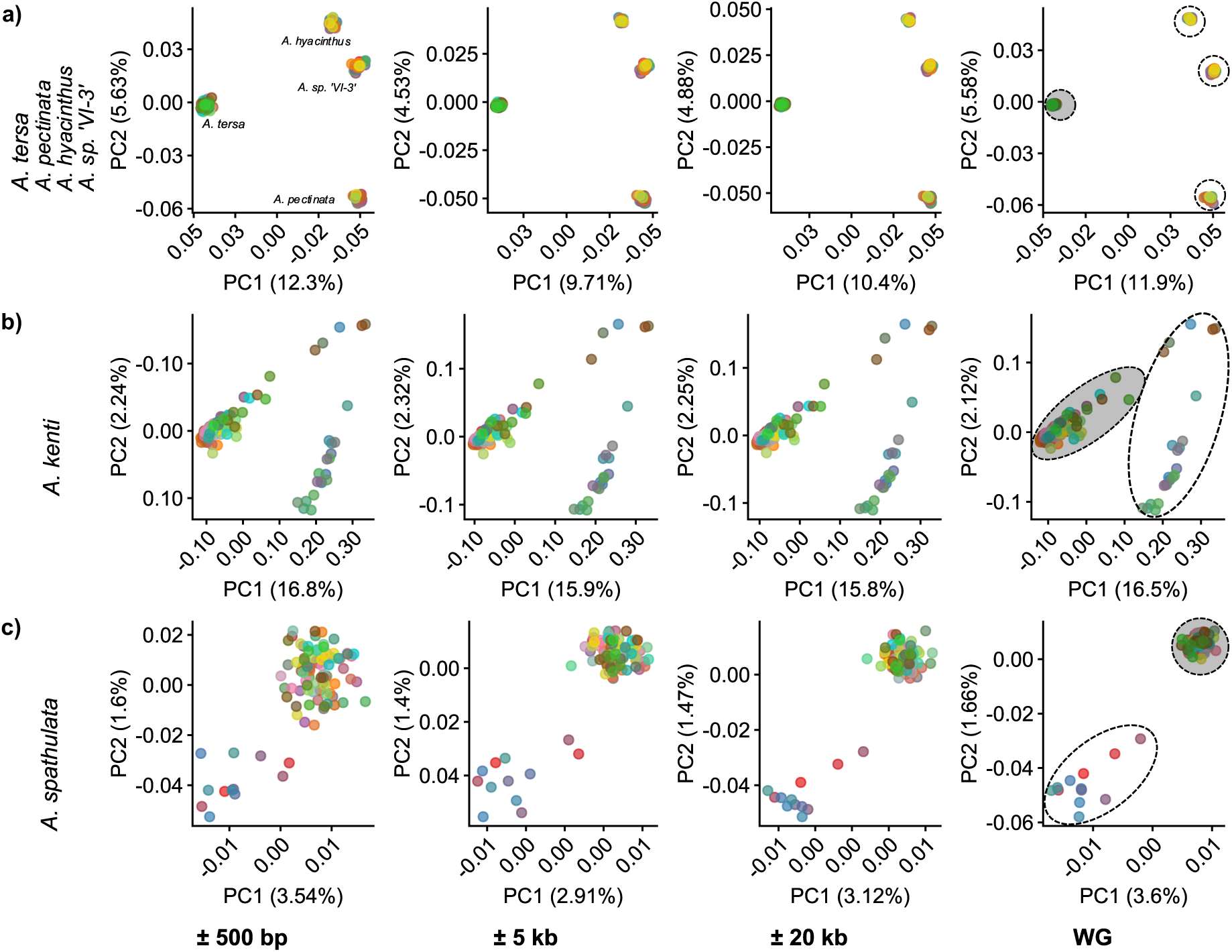
Genomic structure of studied samples in a) *A. tersa, A. pectinata, A. hyacinthus*, *A.* sp. ‘VI-3’, b) *A. kenti* and c) *A.* cf*. spathulata.* The multi-dimensional scaling plots were based on genomic distances calculated from SNPs derived from 1781 UCE loci. Plots illustrate the first two dimensions separating the samples into major clusters (circled; clusters selected for fine-scale analysis shaded in grey). *A. tersa, A. pectinata, A. hyacinthus*, *A.* sp. ‘VI-3’ samples separated into four genetic clusters each representing a distinct species, while *A. kenti* and *A.* cf*. spathulata* samples separated into two major clusters. Samples coloured by conspecific individuals corresponding among panels to allow tracking across extended UCE regions of different lengths.

To ensure intraspecific resolutions, further analyses were applied to the most dominant clusters respectively identified from the MDS (Fig. 2; grey-shaded clusters based on WG). Due to small sample sizes in *A. hyacinthus*, *A.* sp. ‘VI-3’ and *A. pectinata*, *A. tersa* was focussed for the four tabular coral species. In *A. tersa* and *A.* cf. *spathulata*, broad clustering patterns were maintained across extended UCE regions and consistent to the WG benchmark, however at the fine scale, only the *A. kenti* outliers were positioned consistently across extended UCE region lengths and respective to the WG benchmark (Fig.3a, c). In *A. kenti*, broad- and fine-scale genetic diversity clustering were consistent across all extended UCE region sizes and the WG benchmark data, with increasing size showing tighter grouping and finer scale variation (Fig. 3b). Pairwise Mantel tests comparing pairwise genetic distances inferred from the data mapped to each of the extended UCE region references to the WG reference showed a positive correlation across all comparisons (Fig. 4a; *p* = 0.001; Supp. Table 1). In *A. tersa* and *A.* cf*. spathulata*, the correlation coefficients with the WG benchmark increased with extended UCE region size, while still showing moderate correlation at shorter extended UCE regions sizes (UCE ± 500 bp; *r =* 0.617, *r =* 0.473, respectively). In *A. kenti*, *r* values remain consistent across all comparisons (*r* = 0.99).

**Figure 3.**
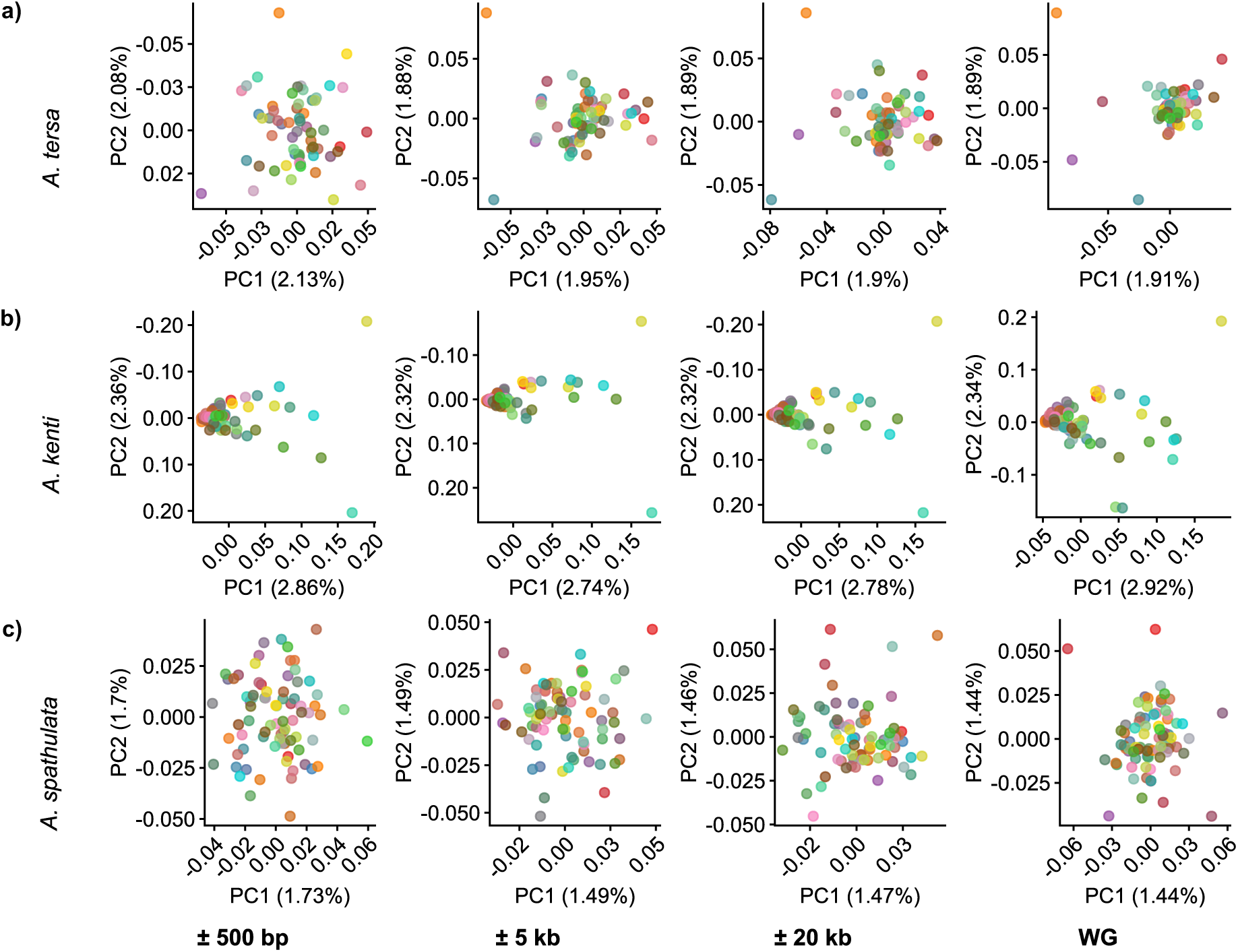
Intraspecific genomic structure in a) *A. tersa*, b) *A. kenti* and c) *A.* cf*. spathulata* from samples belonging to the main clusters identified from WG genomic structure analysis on all conspecific samples. The multi-dimensional scaling plots were based on genomic distances calculated from SNPs derived from 1781 UCE loci. Samples coloured by conspecific individuals corresponding among panels to allow tracking across extended UCE regions of different lengths.

**Figure 4.**
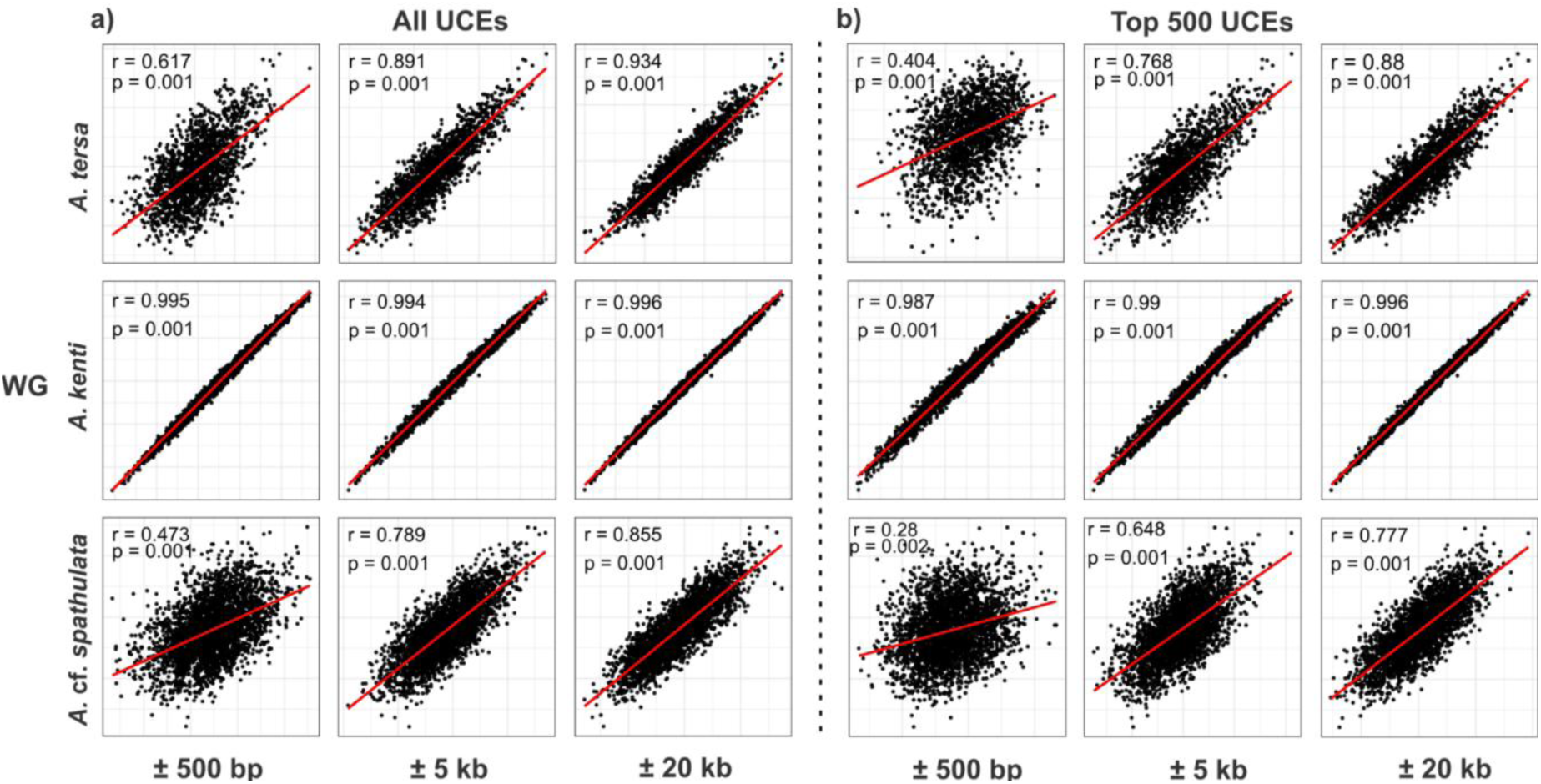
Scatterplots of pairwise comparisons between genetic distance matrices from a) all 1,781 UCE loci or b) top 500 loci with the highest phylogenetic informativeness on the x-axis against that of genome-wide SNPs on the y-axis, with three different lengths of extended UCE regions (± 500 bp, ± 5 kbp and ± 20 kbp).

**Table 1.** Estimates of intraspecific variation calculated from SNP loci captured within extended UCE regions of different window lengths. Data under WG indicate the diversity estimates derived from the whole genome sequencing data. SNPs derived from 1,688 UCE loci in *A. tersa*, 1,715 UCE loci in *A. kenti*, and 1,702 in *A.* cf*. spathulata*. Values are presented as means ± SD.

| Species | Source of SNP loci | $\Pi$ per nucleotide | $\Theta$ per nucleotide |
| --- | --- | --- | --- |
| *A. tersa* | UCE±500 bp | 0.02 ± 0.01 | 0.02 ± 0.01 |
| | UCE±1 kbp | 0.03 ± 0.02 | 0.03 ± 0.02 |
| | UCE±2 kbp | 0.03 ± 0.02 | 0.03 ± 0.02 |
| | UCE±5 kbp | 0.03 ± 0.02 | 0.03 ± 0.02 |
| | UCE±10 kbp | 0.03 ± 0.02 | 0.03 ± 0.02 |
| | UCE±20 kbp | 0.03 ± 0.02 | 0.03 ± 0.02 |
| | WG | 0.02 ± 0.1 | 0.02 ± 0.1 |
| *A. kenti* | UCE±500 bp | 0.02 ± 0.01 | 0.03 ± 0.02 |
| | UCE±1 kbp | 0.03 ± 0.02 | 0.03 ± 0.02 |
| | UCE±2 kbp | 0.03 ± 0.02 | 0.04 ± 0.02 |
| | UCE±5 kbp | 0.03 ± 0.02 | 0.04 ± 0.02 |
| | UCE±10 kbp | 0.02 ± 0.03 | 0.03 ± 0.02 |
| | UCE±20 kbp | 0.02 ± 0.02 | 0.03 ± 0.02 |
| | WG | 0.01 ± 0.04 | 0.02 ± 0.02 |
| *A. cf. spathulata* | UCE±500 bp | 0.02 ± 0.01 | 0.02 ± 0.01 |
| | UCE±1 kbp | 0.02 ± 0.01 | 0.03 ± 0.02 |
| | UCE±2 kbp | 0.03 ± 0.02 | 0.03 ± 0.02 |
| | UCE±5 kbp | 0.03 ± 0.02 | 0.03 ± 0.02 |
| | UCE±10 kbp | 0.03 ± 0.02 | 0.03 ± 0.02 |
| | UCE±20 kbp | 0.03 ± 0.02 | 0.03 ± 0.02 |
| | WG | 0.02 ± 0.02 | 0.02 ± 0.02 |

Estimates of intraspecific genetic variation based on 1,688 UCE-associated extended loci were consistent across all region sizes and the WG benchmark for all three species (Table 1), with polymorphism levels remaining stable regardless of the size of the extended UCE regions or across the genome. In *A. tersa*, mean *π* and *θ* per nucleotide ranged from 0.02 ± 0.01 to 0.03 ± 0.02, indicating high polymorphism among the analysed samples. In *A. kenti*, mean per nucleotide values across all UCE references ranged from 0.03 ± 0.01 to 0.04 ± 0.02 for *π* and 0.02 ± 0.01 to 0.03 ± 0.02 for *θ*, whereas the WG reference produced lower diversity estimates (*π* = 0.01 ± 0.04, *θ* = 0.02 ± 0.02). In *A*. cf. *spathulata*, *π* and *θ* estimates from 1,702 UCE loci ranged from 0.02 ± 0.02 to 0.03 ± 0.02 for both statistics.

### Analysis on UCE loci subset

Resolving phylogeny and assessing intraspecific variation from a reduced set of loci were examined using three subsets of 500, 200 and 100 loci out of the 1,781 UCEs, selected based on the highest number of parsimony informative sites. When performing ML inference on the 500 UCE subset, with 433 UCEs meeting the >75% sample coverage threshold, 54.7% of the internal nodes were resolved with IQ-TREE bootstrap support values of 100 (70.5% with ≥ 95 bootstrap support) (Fig. 5). The tree reconstructed the six *Acropora* clades with 100% bootstrap support, in which each *Acropora* species formed a monophyletic lineage (≥95% bootstrap). In contrast, *A. kenti* monophyly was not reconstructed in the 200 and 100 loci trees. On the 200 loci tree, 173 loci were present in the 75% completeness matrix, and 46% of nodes were resolved IQ-TREE bootstrap values of 100 (59.7% ≥ 95) (Supp. Fig. 2). The six-clade structure was reconstructed with 100% bootstrap support for each clade, however, the *A. kenti* samples were dispersed throughout the clade I. For the 100 UCE subset, 84 loci were present in the 75% completeness matrix. In the ML tree, 41.7% of the internal nodes were resolved with IQ-TREE bootstrap values of 100 (61.9% ≥ 95; Supp. Fig. 3). The six-clade structure was preserved with good support (100 bootstrap value for each clade), however the monophyly of *A. pectinata, A.* sp. ‘VI-3’, and *A. kenti* was not reconstructed. *A.* cf*. spathulata*, *A. tersa*, and *A. hyacinthus* remained reciprocally monophyletic. The results indicate well supported clades and fit of the *A. tersa* (98 bootstrap value), *A. hyacinthus* (100 bootstrap value), and *A.* cf*. spathulata* samples (100 bootstrap value). The *A. kenti* samples were dispersed throughout clade I and did not form a monophyletic group (77 bootstrap value).

**Figure 5.**
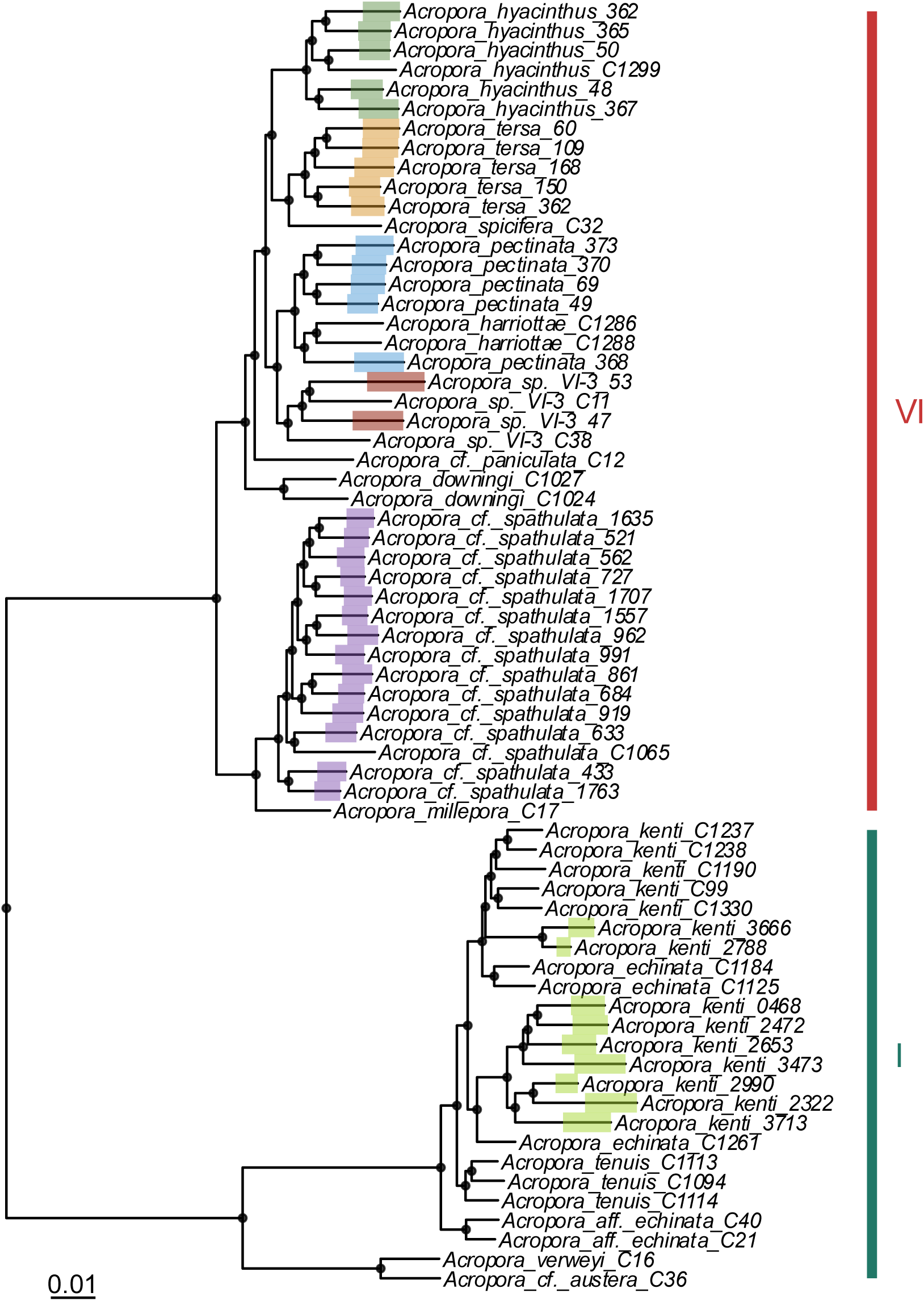
Maximum likelihood tree based on edge trimmed alignments of 433 UCE loci (default UCE length, core ± ∼1kbp). Samples of each of the 6 species highlighted by branch colour, and reference samples from Cowman et al. (2020) with no highlight. Dots indicate nodes with bootstrap support ≥ 95. Vertical bars indicate clades VI and I.

MDS analysis on the 500 loci subset showed consistent clustering patterns to the all UCE-loci and WG mapped data, with finer scale resolution and tighter clustering at increasing extended UCE region size and loci number (Supp. Fig. 4). On the 200 and 100 loci datasets (Supp. Fig. 6-7), clustering was broadly similar to the WG-based MDS, with lower resolution across the reference extended UCE region sizes. *A. kenti* showed high correlation across all extended UCE region sizes (r = 0.99, p = 0.001; Fig. 4b). Mantel tests of pairwise genetic distance in *A*. cf*. spathulata* and *A. tersa* indicate low correlation with the WG data at short extended UCE region sizes (± 500 bp; r = 0.28 and r = 0.404, respectively; p = 0.001) and high correlation at long extended UCE region sizes (± 20 kbp; r = 0.777 and r = 0.88, respectively p = 0.001). Although clustering was similar to the WG MDS, the 200 and 100 loci subsets showed very low and non-significant correlation to the WG benchmark across all extended UCE region lengths (r = 0.02-0.66, p>0.001; Supp. Fig. 5-6).

Estimates of *π* and *θ* per nucleotide on the 500 loci dataset followed a similar pattern to those observed in the all-loci dataset in two species (Supp. Table 2). In *A. tersa*, *π* and *θ* varied between 0.01 ± 0.1 and 0.03 ± 0.07 across all reference sizes and were consistent with the WG-derived values (0.02 ± 0.01). In *A.* cf*. spathulata*, estimates remained stable across all extended UCE region sizes and the WG reference, with *π* values of 0.03 ± 0.01 - 0.05 ± 0.02 and *θ* per locus values of between 0.03 ± 0.01 - 0.04 ± 0.02. However, in *A. kenti*, both estimates of diversity were only consistent to the WG benchmark at the ± 500 bp, ± 1 kbp, and ± 2 kbp extended UCE regions (0.02 ± 0.01 - 0.04 ± 0.01) while increased at the ± 5 kbp, ± 10 kbp and ± 20 kbp references (0.04 ± 0.01 - 0.09 ± 0.05). In the 200 and 100 loci subsets, all estimates of intraspecific variation varied between 0.02 ± 0.01 and 0.04 ± 0.02 and were consistent across all sources of SNP loci (UCE-derived and WG-derived) (Supp. Tables 3-4).

## Discussion

This study tested the feasibility of using extended UCE regions to study interspecific and intraspecific genetic variation of coral taxa. Using population wide genomic data and a reduced loci approach, the results indicate that extended UCE regions have the potential to be used for fast and scalable genetic assessment at both interspecific and intraspecific scales. This supports streamlined genetic screening workflows where fast and scalable genetic assessment are valuable, such as during coral spawning associated aquaculture reproduction. Within the limited time windows available for spawning and progeny production, such approach enables timely species identification of broodstock corals and assessment of offspring genetic diversity in support of reef restoration programs.

### Taxonomic verification established based on UCE-associated flanking regions

The placement of *A*. *tersa*, *A. hyacinthus*, *A.* sp. ‘VI-3’, *A. pectinata*, *A. kenti*, and *A.* cf*. spathulata* samples based on 1,177 UCE-associated loci confirmed their taxonomic expectations from the previous studies (Cowman et al., 2020; Rassmussen et al., 2025), forming respective clades including reference sequences within the six-clade structure in the *Acropora* phylogeny (Cowman et al., 2020). Moreover, this study confirms that integration of lcWGS to a UCE targeted phylogeny provides a practical approach to resolved species identification with molecular evidence. For example, field collection using the characters used to identify *A. hyacinthu*s under traditional morphological taxonomy encompassed at least four distinct, sympatric species on the Great Barrier Reef (Rassmussen et al., 2025; Naugle et al. 2024). In population genetic analyses, inadvertently pooling individuals from distinct species inflates genetic diversity estimates and obscures true population structure, thus conclusions about diversity, connectivity, and population structure may reflect an artificial composite of two species rather than either one accurately (Sheets et al., 2018; Warner et al., 2015). Molecular-based verification using this study’s approach can therefore be a practical important step to establish a study basis depending on the scope.

When analysing the 500 UCE subset with the standard UCE lengths (core ± ∼1kbp), all species are resolved and placed correctly in the six-clade tree (based on 433 out of the 500 UCE loci that passed the sample coverage cutoff). This study indicates that the efficiency of the species identification can be improved by using a subset of UCE-associated loci. However, when using the reduced 100 and 200 loci set (173 and 84 loci post filtering, respectively), while the placement within clades I and VI was consistent and well supported, species-level patterns were not always retained. Specifically, *A. pectinata, A.* sp. ‘VI-3’, and *A. kenti* samples did not form respective monophyletic groups. ML inference is generally performed on thousands of genome-wide loci (Cowman et al., 2020; Hughes et al., 2018), while studies have characterised loci subsets that can generate taxonomically informative SNPs and delineate coral species effectively (Mongiardino Koch 2021, Ramírez-Portilla et al., 2022). For example, SNPs called from 79 UCE loci were able to differentiate three closely related species in the Clade VI (*A.* cf*. bifurcata, A.* cf*. cytherea* and *A. hyacinthus*).

The taxa-specific loss of phylogenetic resolution when reducing the number of loci from 1,177 to 84 is likely attributed to the specific subset of UCE loci being more informative to better resolve certain taxa than other. In our study, sub-setting UCE-loci was based on parsimony informativeness throughout the studied taxa encompassing the genus *Acropora*. When targeting a specific range of species in the tree, a minimum set of UCE loci to differentiate them should reflect the target taxonomic range. Thus, the methodology for loci selection can be improved by incorporating prior screening of loci that display allelic exclusivity for each species or genetic cluster, which can be tailored to the range of taxa of interest (Ramírez-Portilla et al., 2022).

### Intraspecific genetic variation is captured in extended flanking regions of UCE loci with comparable accuracy to a whole genome benchmark

SNPs captured from the full set of extended UCE-associated regions can resolve intraspecific genetic variation with comparable accuracy to genome-wide SNPs across lengths of flanking regions associated with UCE loci studied. Our results highlight that the size of flanking regions and the number of associated UCE loci can be modulated to further improve the data-efficiency without losing valuable genetic information.

Visualization of population genetic structuring using MDS based on genetic distances from the all-loci dataset reproduced the consistent clustering patterns using the WG approach regardless of the extended UCE region sizes (Fig. 2). Consistent to the above ML phylogeny and the previous genomic and morphological evidence (Rassmussen et al., 2025; Naugle et al., 2024), the MDS analyses encompassing the *A*. *tersa*, *A. hyacinthus*, *A.* sp. ‘VI-3’, and *A. pectinata* samples reveals four distinctive clusters that delineates four distinct species. The clusters in *A. kenti* correspond to distinct populations that are likely exchanging low levels of geneflow, as previously reported by Matias et al. (2023) potentially reflecting genetically distinct taxa; the shaded cluster corresponds to a widespread taxon, whereas the smaller cluster consists of individuals sampled from northern offshore reefs below 14.7 latitude. Similarly, the two groups in the *A.* cf*. spathulata* samples correspond to distinct populations, with the tighter cluster corresponding to samples from the Capricorn-Bunker group in the southern GBR and the rest from the central and northern GBR. This region often shows genetic differentiation from populations on the rest of the GBR due to restricted gene flow by geographical isolation (Matias et al., 2023; Van Oppen et al., 2011, 2015) to the extent that the region may harbour numerous distinct, endemic species. Two such species (*Acropora harriottae* Baird & Rassmussen, 2025 and *Cyphastrea salae* Baird et al., 2017) have been described to date, although many more are known and awaiting description. These findings further emphasise the problems with assuming a species is widespread across the GBR and highlight the need to quantitatively assess genetic structure within putatively widespread ‘species’.

The genetic distance matrices produced by the 500 UCE subset and the WG benchmark indicate low positive correlation at short extended UCE region lengths in *A. tersa* and *A.* cf. *spathulata*, however the correlation strengthens above the ± 5 kbp extended UCE region. Increasing length after ± 5 kbp does not improve resolution or correlation with the WG dataset when analysing the all-loci dataset, but it does increase resolution on the reduced loci subset. Similarly, the genetic distance matrices showed strong positive correlation with the genome-wide data across the all-loci extended UCE references although there is no substantial increase in the strength of the correlation beyond 5 kbp. Correlation between the 100 and 200 loci dataset and the WG data was low and only significant on the 20 kbp extended UCE region.

MDS on the main clusters identified from the full-loci MDS on the 500 loci set shows the same clustering patterns as the WG approach, with better resolution and finer-scale variation visible at extended UCE regions ± 5, 10 and 20 kbp. When increasing to 500 loci, resolution is kept at the ± 20 kbp extended UCE region and correlates strongly with the WG results. Future work should thus incorporate a subset of at least 500 UCEs to be able to identify fine-scale genetic structure. However, resolution and fine-scale patterns are lost in all species when loci are further reduced, with 100 and 200 loci not being enough to differentiate the main genetic groups (Supp. Figs. 5-6). Reducing loci in populations with closely related individuals such as *A.* cf*. spathulata* and *A. tersa* likely obscures fine scale differentiation; this relatedness also drives the low correlation with the all-loci dataset. Previous studies in other *Acropora* species reported high polymorphism in *A. cervicornis* from Florida (Hemond & Vollmer, 2010), *A.* cf. *tenuis* from Japan (Zayasu et al., 2018) and *A. millepora* and *A. kenti* from the GBR (Matias et al., 2023; Van Oppen et al., 2011). Estimates of *π* in *A. kenti* reported in (Matias et al., 2023), were similar to this study (*π =* 0.02). *θ* estimates were closer to the genome-wide approach in this work (*θ* = 0.02-0.03), but not to the extended UCE region approach (*θ* = 0.04-0.05). The genetic diversity and structure in *Acropora* is potentially due to the geneflow and connectivity driven by the broadcast spawning sexual reproduction strategy of the group (Van Der Ven, Heynderickx, et al., 2021).

When using 500, 200 and 100 UCE loci to examine estimates of genetic variation, UCE-derived SNP loci at all ∼1,700 extended UCE region lengths produced results consistent with those from WG-derived SNP loci in *A. tersa* and *A.* cf*. spathulata*. However, when using the 500 loci subset on *A. kenti*, the estimates increase significantly with extended UCE region size, while this was not evident in all UCE loci or the subsets of 100 and 200. When selecting highly informative loci for genomic analysis, extended regions of certain loci may contain high genetic variability among samples resulting in a SNP ascertainment bias, population differentiation can be overestimated (high-grading bias; Lachance & Tishkoff, 2013; Lee et al., 2025). Additionally, low read depth can lead to difficulties distinguishing between heterozygotes and homozygotes. In the case of the current study, SNP ascertainment, low-quality shallow lcWGS and the large genetic variation between samples specific to the *A. kenti* samples, are likely overestimating the diversity estimates with increasing extended UCE region size. The values from the extended UCE-loci references are consistent between each other, regardless of the higher SNP count associated with the size of the reference.

Taken together, it is recommended to extend the area around the UCE core and flanking regions to at least 5 kbp either side, resulting in sequences approximately 10 kbp long which align with long-read sequencing platforms where 10-100 kbp reads are commonly processed (Amarasinghe et al., 2020). Pairing this with a subset of 500 informative loci opens the potential to improve the efficiency of the sequencing effort to characterize genetic variation of coral populations.

### Extended UCE regions as target for a promising approach to rapid genetic assessment: Broader implications and outlook

This study demonstrates that extended UCE regions are a practical dual-purpose tool for both species identification and profiling of intraspecific genetic variation. This approach has the potential to make the sequencing effort more efficient and to be applied to a long-read sequencing platform. A region of at least 5 kbp upstream and downstream of each UCE locus provides a good balance between plateaued resolution and sequencing effort, thus making it a useful target for a fast and scalable workflow. Furthermore, a subset of 500 informative UCE-associated regions showed enough resolution to retain both phylogenetic and intraspecific differentiations, offering a data- and cost-effective alternative to WG sequencing or target capture of all ∼1,700 UCE loci.

The ability to accurately identify and characterize the genetic makeup of populations is fundamental for effective management and conservation strategies. In coral reef studies and emerging coral restoration endeavours, precise and rapid species identification is critical for biological studies for contextualising physiological studies, ecological observations and behavioural patterns (Guedes et al., 2025). Similarly, in population genetics, the misidentified or cryptic species can obscure and bias interpretation of data leading to erroneous conclusions regarding population structure, gene flow or physiological tolerance within a species (Gómez-Corrales & Prada, 2020; Sheets et al., 2018). By resolving these taxonomic issues based on multi-locus SNP data, this approach can support more accurate, robust and potentially more efficient genetic assessments across taxa and evolutionary scales. This work lays the groundwork for the development of a scalable assay for coral species identification and assessment of genetic variation of coral populations. In restoration aquaculture, for example, a rapid scalable assay will allow timely parental selection and broodstock quality control checks to ensure that the sexual propagation of coral populations captures sufficient genetic diversity in offspring for responsible and effective coral restoration efforts that deploy captive bred corals. The novel methodology developed in this study also presents a foundation to robust and practical approach for genetic assessment for other traditionally problematic taxa in a short timeframe to underpin findings of experimental studies with genetic characterisation.

A long-region specific capturing and enrichment would be a required next step to improve genotyping accuracy by increasing sequence depths per effort. The region specific enrichment method produces fragments of >20 kbp with higher coverage of target regions (Dapprich et al., 2016). Furthermore, speed of genetic data acquirement can be greatly improved by incorporating an on-demand, portable long-read sequencing platform such as Oxford Nanopore Technologies sequencers (Wang et al., 2021). In addition, further *in-silico* and *in-vitro* studies to improve loci prioritisation and sub-setting for taxonomic verification and intraspecific genetic profiling can improve the scalability and effectiveness of this assay.

## Author contributions

Conceptualisation: Y.S.; Methodology: A.M with input from Y.S. and P.C.; Analysis: A.M. with input from Y.S. and P.C.; Writing – original draft: A.M.; Writing – review and editing: A.M., Y.S., P.C., T.B., Y.Y., D.B.

## Acknowledgements

This work was undertaken in the Reef Restoration and Adaptation Program (RRAP), funded by the partnership between the Australian Government’s Reef Trust and the Great Barrier Reef Foundation and partners including the Australian Institute of Marine Science, and James Cook University. We thank Prof. Cynthia Riginos and Dr. Lorenzo Bertola for valuable discussion for this study. We thank Dr. Iva Popovic, Dr. Emily Howells, Dr. Katharine Prata, and Dr. Hugo Denis for providing us access to the genomic data used in this study. P.F.C and T.B were also funded by ARC DECRA Fellowships (DE170100516 and DE180100746, respectively) and the Queensland Museum’s Project DIG.

## Data Availability and Benefit-Sharing

Scripts, metadata and list of candidate UCE loci are available at: https://github.com/amateos1998/Extended-UCE-regions-for-genetic-assesment

## SUPPLEMENTARY FIGURES

**SF1.**
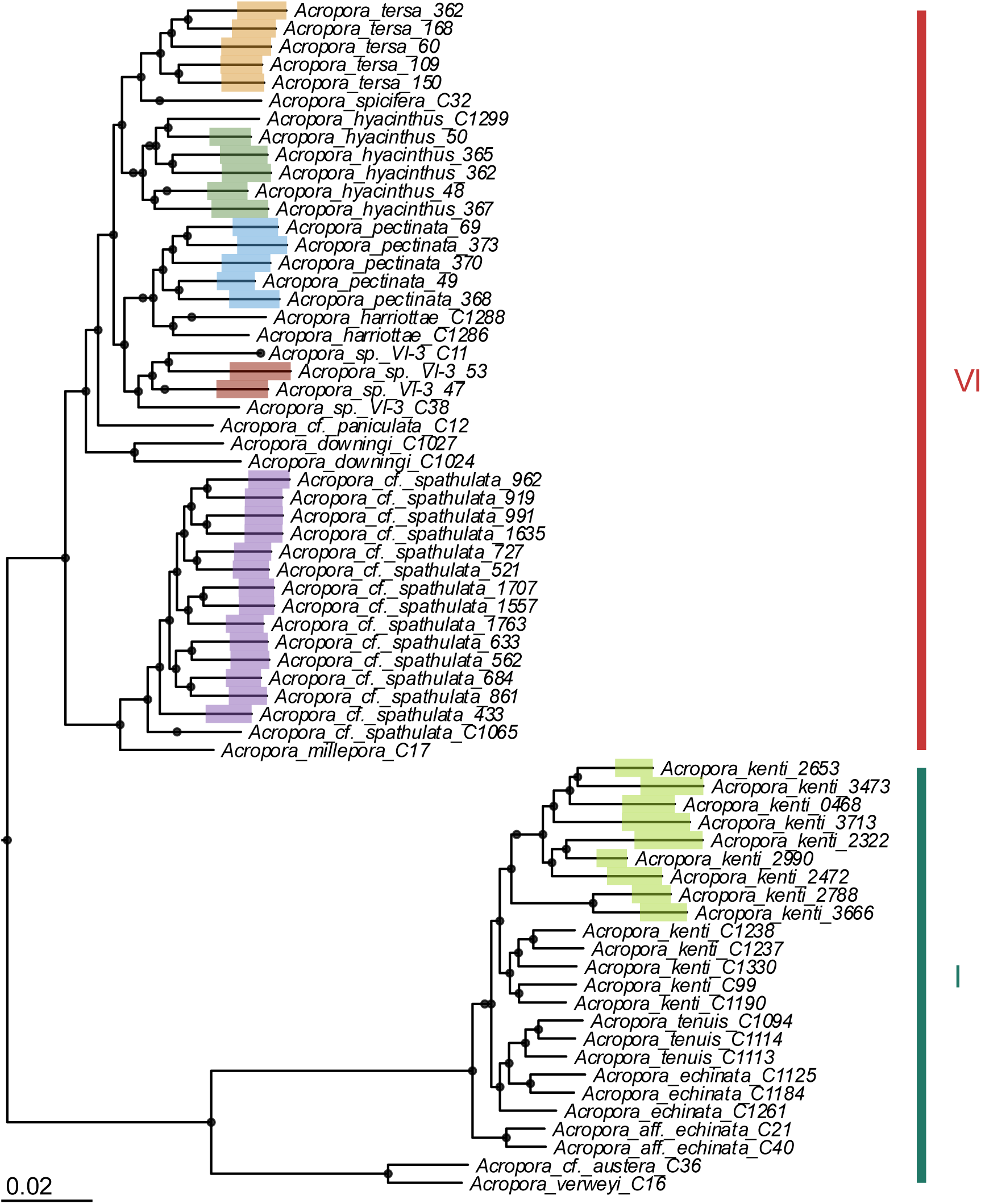
Maximum likelihood tree (0.99 bootstrap value) based on edge trimmed alignments of 1,177 UCE loci (default UCE length, core ± ∼1kbp). Dots indicate nodes with bootstrap support ≥ 95. Samples of each of the 6 species highlighted by colour, Cowman et al. 2020 samples with no highlight. *Isopora* cf. *brueggemanni* used as outgroup. Vertical bars indicate clades VI and I.

**SF2.**
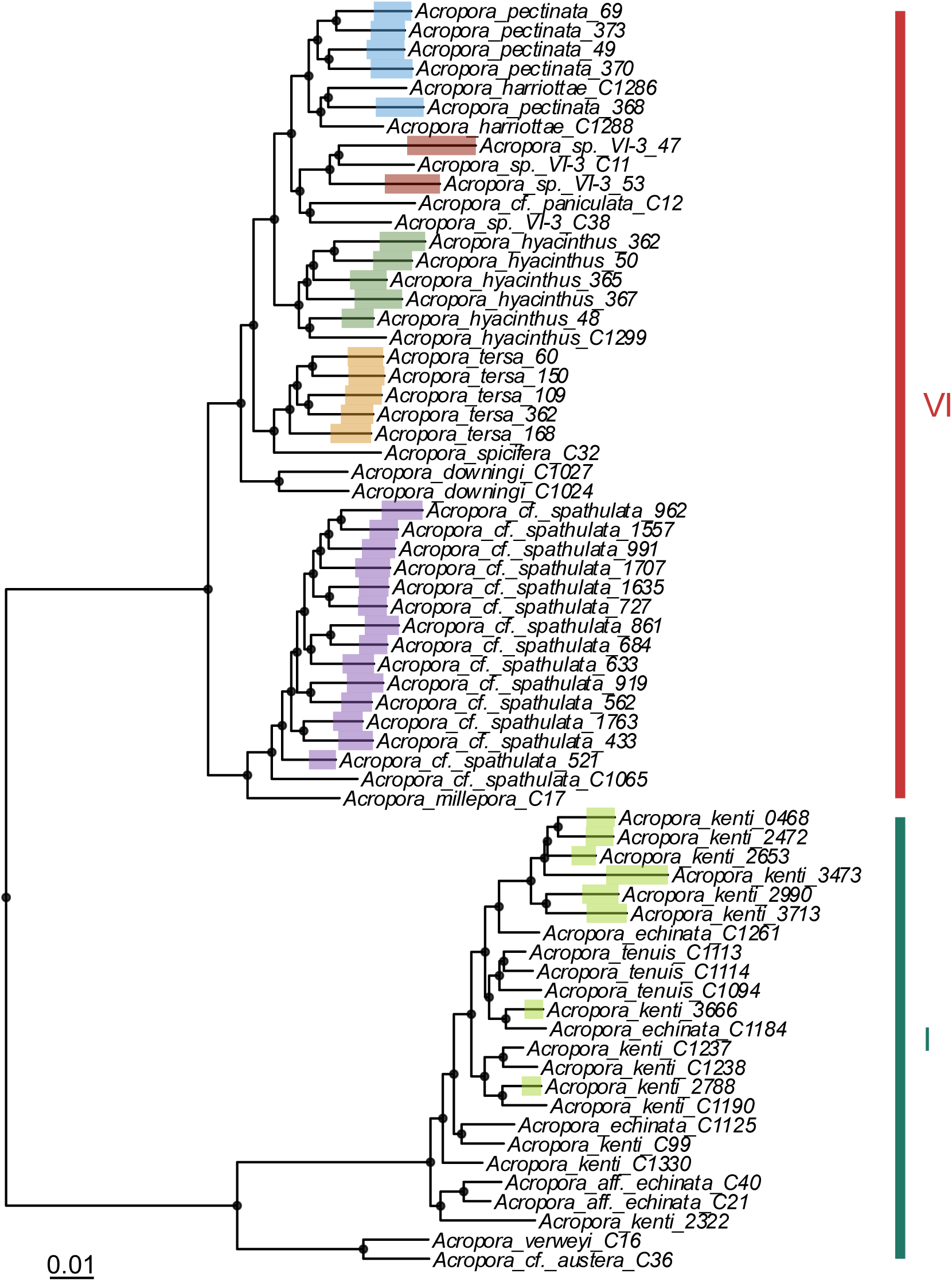
Maximum likelihood tree (0.99 bootstrap value) based on edge trimmed alignments of 173 UCE loci (default UCE length, core ± ∼1kbp). Dots indicate nodes with bootstrap support ≥ 95. Samples of each of the 6 species highlighted by colour, Cowman et al. 2020 samples with no highlight. *Isopora* cf. *brueggemanni* used as outgroup. Vertical bars indicate clades VI and I.

**SF3.**
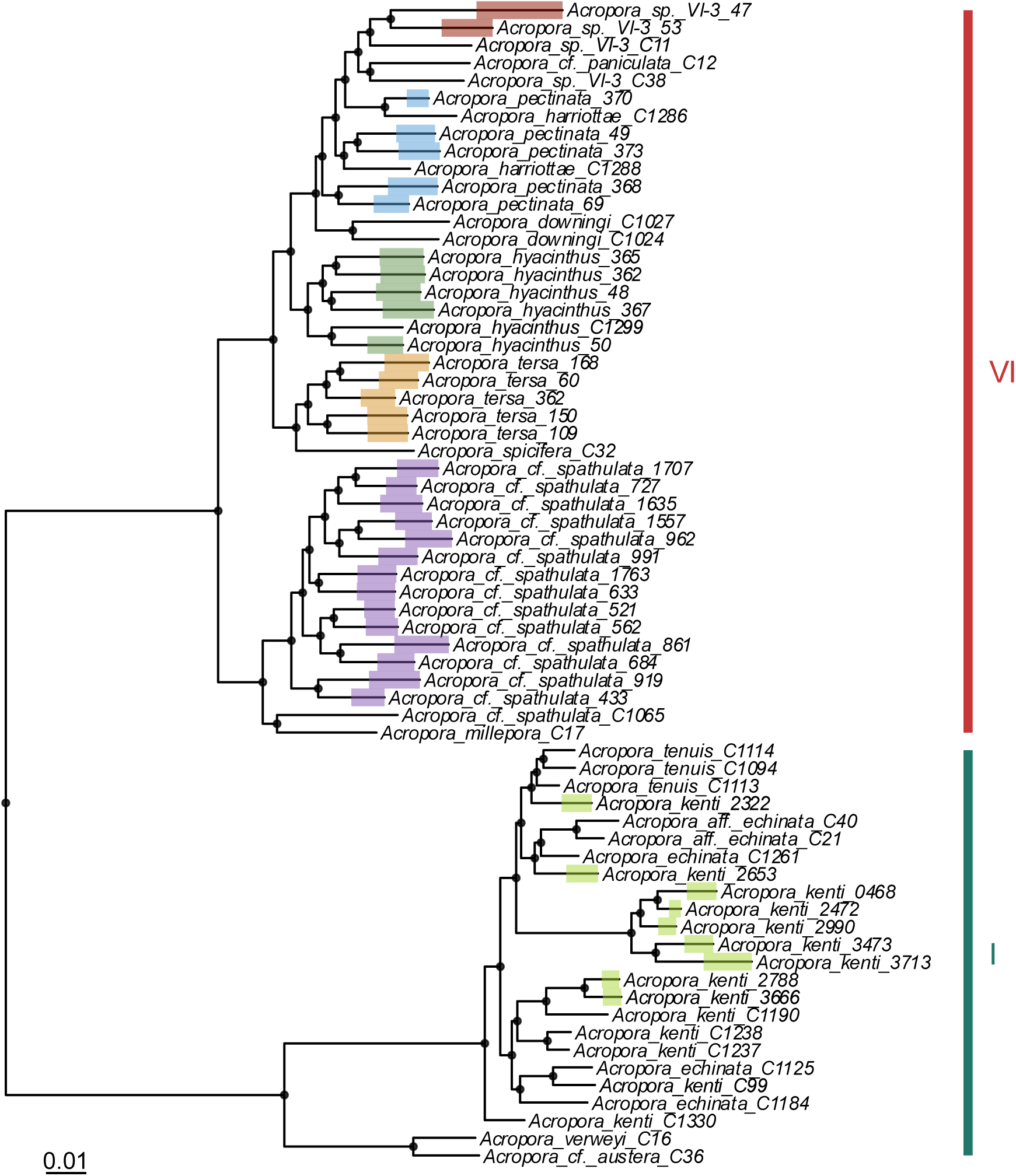
Maximum likelihood tree (0.99 bootstrap value) based on edge trimmed alignments of 84 UCE loci (default UCE length, core ± ∼1kbp). Dots indicate nodes with bootstrap support ≥ 95. Samples of each of the 6 species highlighted by colour, Cowman et al. 2020 samples with no highlight. *Isopora* cf. *brueggemanni* used as outgroup. Vertical bars indicate clades VI and I.

**SF4.**
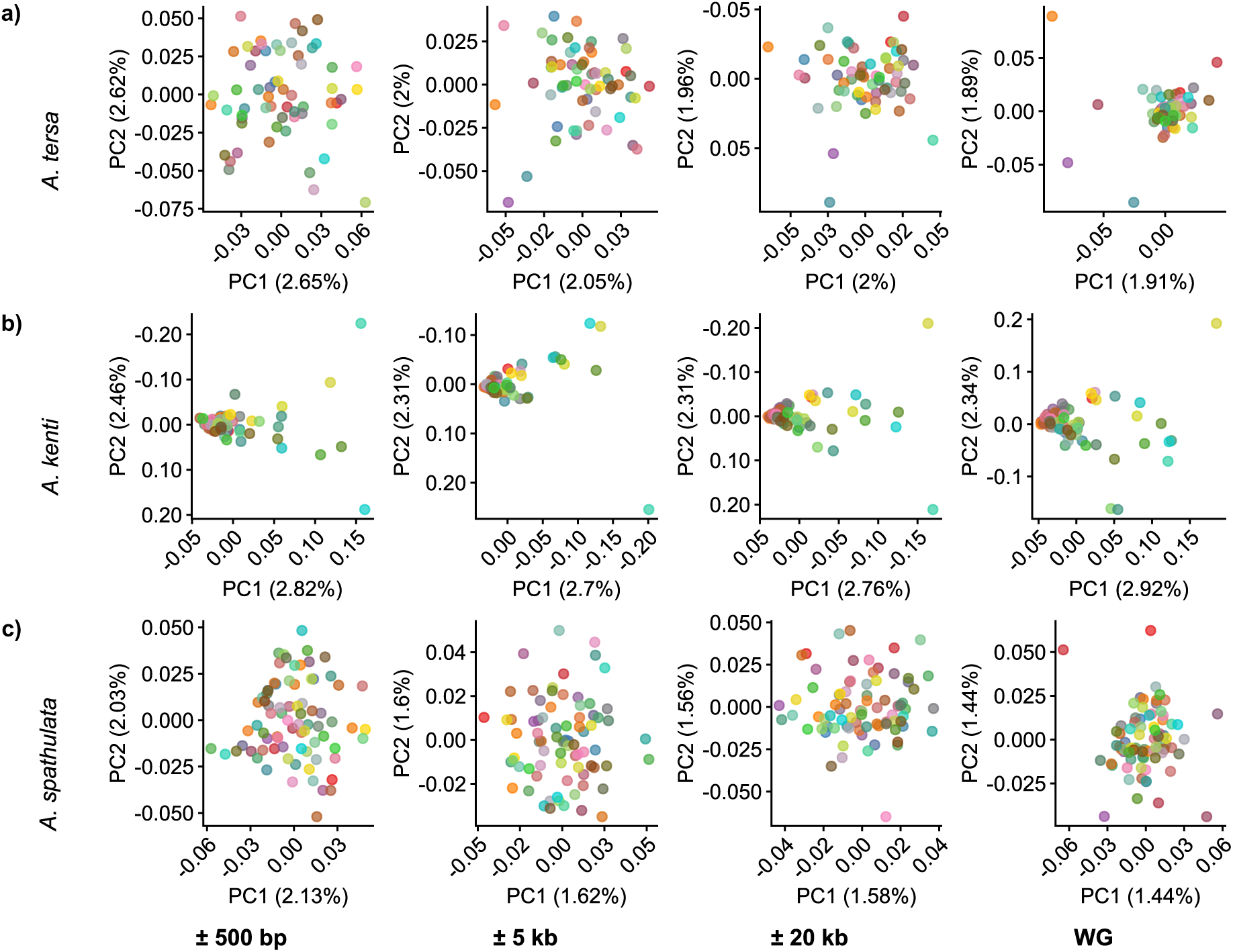
Fine-scale intraspecific genomic structure in a) *A. tersa*, b) *A. kenti* and c) *A.* cf*. spathulata* from main clusters identified from WG MDS analysis on all samples. Genetic distances calculated from SNPs derived from 500 UCE loci. Samples coloured by individual to allow tracking across UCE references of different lengths.

**SF5.**
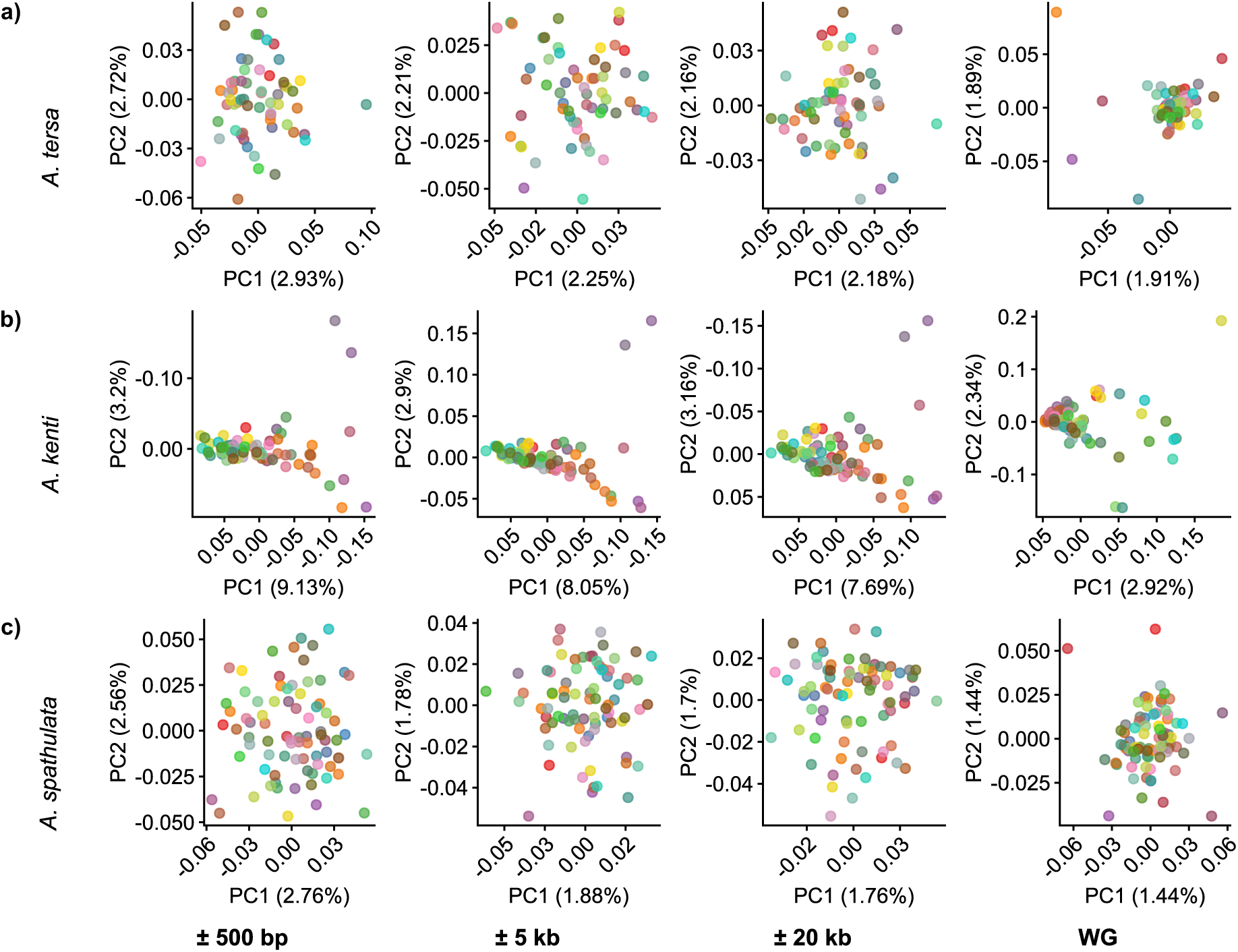
Fine-scale intraspecific genomic structure in a) *A. tersa*, b) *A. kenti* and c) *A.* cf*. spathulata* from main clusters identified from WG MDS analysis on all samples. Genetic distances calculated from SNPs derived from 200 UCE loci. Samples coloured by individual to allow tracking across UCE references of different lengths.

**SF6.**
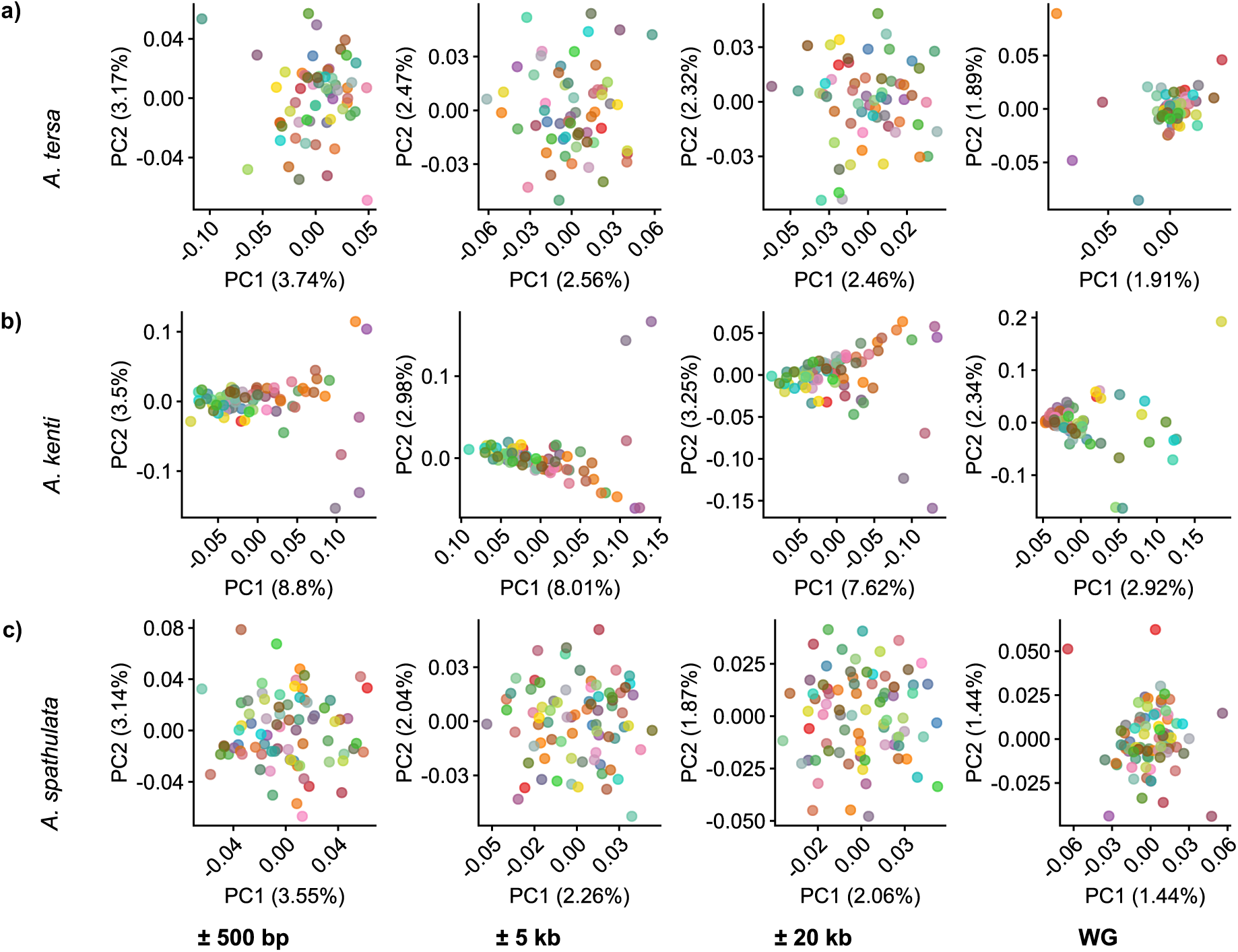
Fine-scale intraspecific genomic structure in a) *A. tersa*, b) *A. kenti* and c) *A.* cf*. spathulata* from main clusters identified from WG MDS analysis on all samples. Genetic distances calculated from SNPs derived from 100 UCE loci. Samples coloured by individual to allow tracking across UCE references of different lengths.

## SUPPLEMENTARY TABLES

**ST1.**
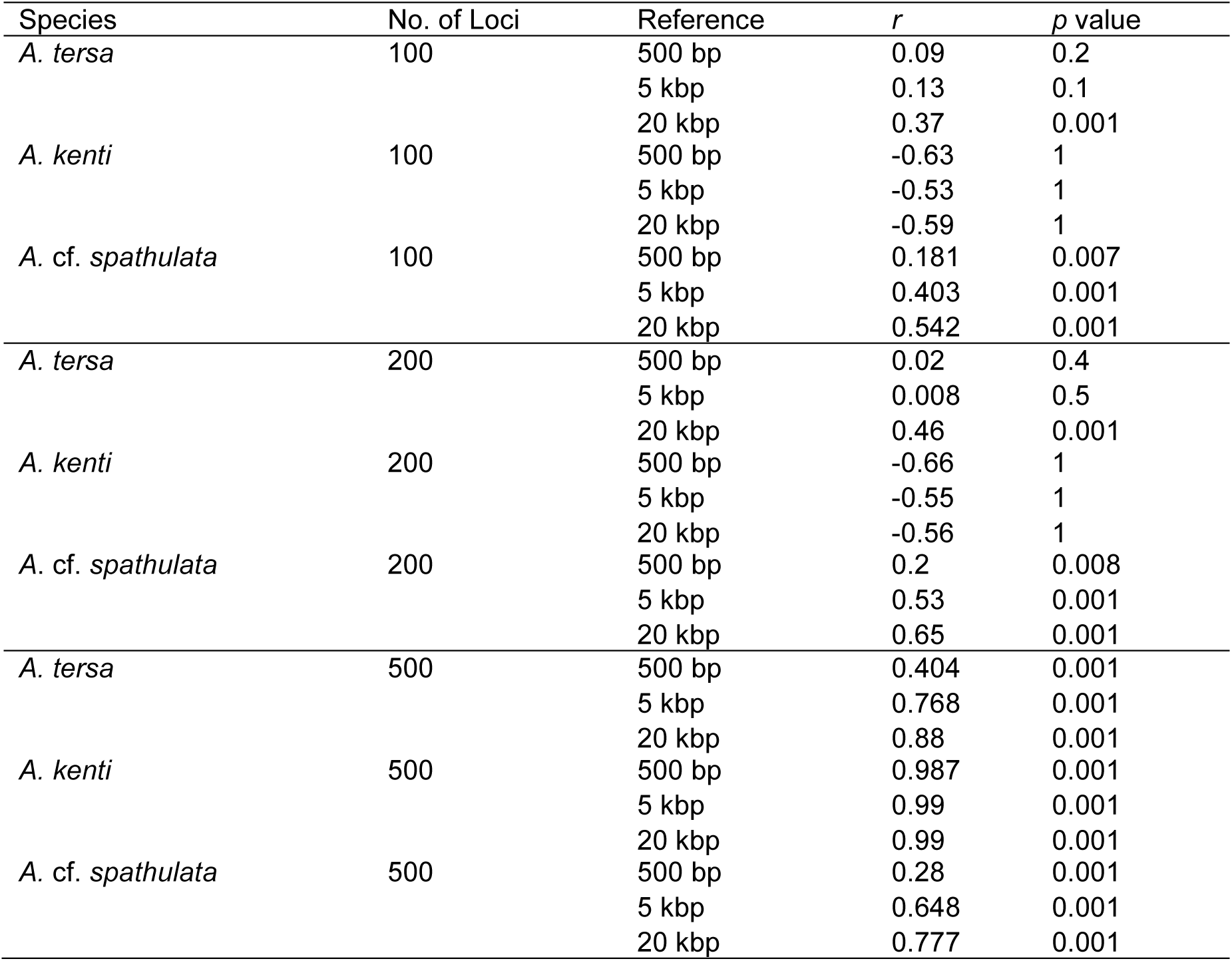
Mantel test results of pairwise comparisons between genetic distance matrices from SNP loci within extended UCE regions and genome-wide SNP loci.

| Species | No. of Loci | Reference | *r* | *p* value |
| --- | --- | --- | --- | --- |
| *A. tersa* | 100 | 500 bp | 0.09 | 0.2 |
|  |  | 5 kbp | 0.13 | 0.1 |
|  |  | 20 kbp | 0.37 | 0.001 |
| *A. kenti* | 100 | 500 bp | -0.63 | 1 |
|  |  | 5 kbp | -0.53 | 1 |
|  |  | 20 kbp | -0.59 | 1 |
| *A. cf. spathulata* | 100 | 500 bp | 0.181 | 0.007 |
|  |  | 5 kbp | 0.403 | 0.001 |
|  |  | 20 kbp | 0.542 | 0.001 |
| *A. tersa* | 200 | 500 bp | 0.02 | 0.4 |
|  |  | 5 kbp | 0.008 | 0.5 |
|  |  | 20 kbp | 0.46 | 0.001 |
| *A. kenti* | 200 | 500 bp | -0.66 | 1 |
|  |  | 5 kbp | -0.55 | 1 |
|  |  | 20 kbp | -0.56 | 1 |
| *A. cf. spathulata* | 200 | 500 bp | 0.2 | 0.008 |
|  |  | 5 kbp | 0.53 | 0.001 |
|  |  | 20 kbp | 0.65 | 0.001 |
| *A. tersa* | 500 | 500 bp | 0.404 | 0.001 |
|  |  | 5 kbp | 0.768 | 0.001 |
|  |  | 20 kbp | 0.88 | 0.001 |
| *A. kenti* | 500 | 500 bp | 0.987 | 0.001 |
|  |  | 5 kbp | 0.99 | 0.001 |
|  |  | 20 kbp | 0.99 | 0.001 |
| *A. cf. spathulata* | 500 | 500 bp | 0.28 | 0.001 |
|  |  | 5 kbp | 0.648 | 0.001 |
|  |  | 20 kbp | 0.777 | 0.001 |

**ST2.** Measures of intraspecific variation calculated from SNP loci captured within extended UCE regions. SNPs derived from 500 UCE loci present on all taxa. Data are presented as means ± SD.

| Species | Source of SNP loci | $\Pi$ per nucleotide | $\Theta$ per nucleotide |
| --- | --- | --- | --- |
| *A. tersa* | UCE±500 bp | 0.01 ± 0.01 | 0.02 ± 0.01 |
| | UCE±1 kbp | 0.01 ± 0.01 | 0.02 ± 0.01 |
| | UCE±2 kbp | 0.03 ± 0.01 | 0.03 ± 0.01 |
| | UCE±5 kbp | 0.03 ± 0.02 | 0.03 ± 0.02 |
| | UCE±10 kbp | 0.02 ± 0.07 | 0.03 ± 0.07 |
| | UCE±20 kbp | 0.01 ± 0.07 | 0.01 ± 0.07 |
| | WG | 0.02 ± 0.1 | 0.02 ± 0.1 |
| *A. kenti* | UCE±500 bp | 0.02 ± 0.01 | 0.03 ± 0.01 |
| | UCE±1 kbp | 0.03 ± 0.01 | 0.04 ± 0.02 |
| | UCE±2 kbp | 0.03 ± 0.01 | 0.04 ± 0.02 |
| | UCE±5 kbp | 0.04 ± 0.01 | 0.04 ± 0.01 |
| | UCE±10 kbp | 0.06 ± 0.02 | 0.06 ± 0.02 |
| | UCE±20 kbp | 0.10 ± 0.05 | 0.09 ± 0.05 |
| | WG | 0.01 ± 0.04 | 0.02 ± 0.02 |
| *A. cf. spathulata* | UCE±500 bp | 0.03 ± 0.01 | 0.03 ± 0.01 |
| | UCE±1 kbp | 0.04 ± 0.01 | 0.03 ± 0.01 |
| | UCE±2 kbp | 0.05 ± 0.01 | 0.04 ± 0.02 |
| | UCE±5 kbp | 0.05 ± 0.01 | 0.04 ± 0.01 |
| | UCE±10 kbp | 0.03 ± 0.01 | 0.03 ± 0.01 |
| | UCE±20 kbp | 0.03 ± 0.01 | 0.03 ± 0.01 |
| | WG | 0.02 ± 0.02 | 0.02 ± 0.02 |

**ST3.**
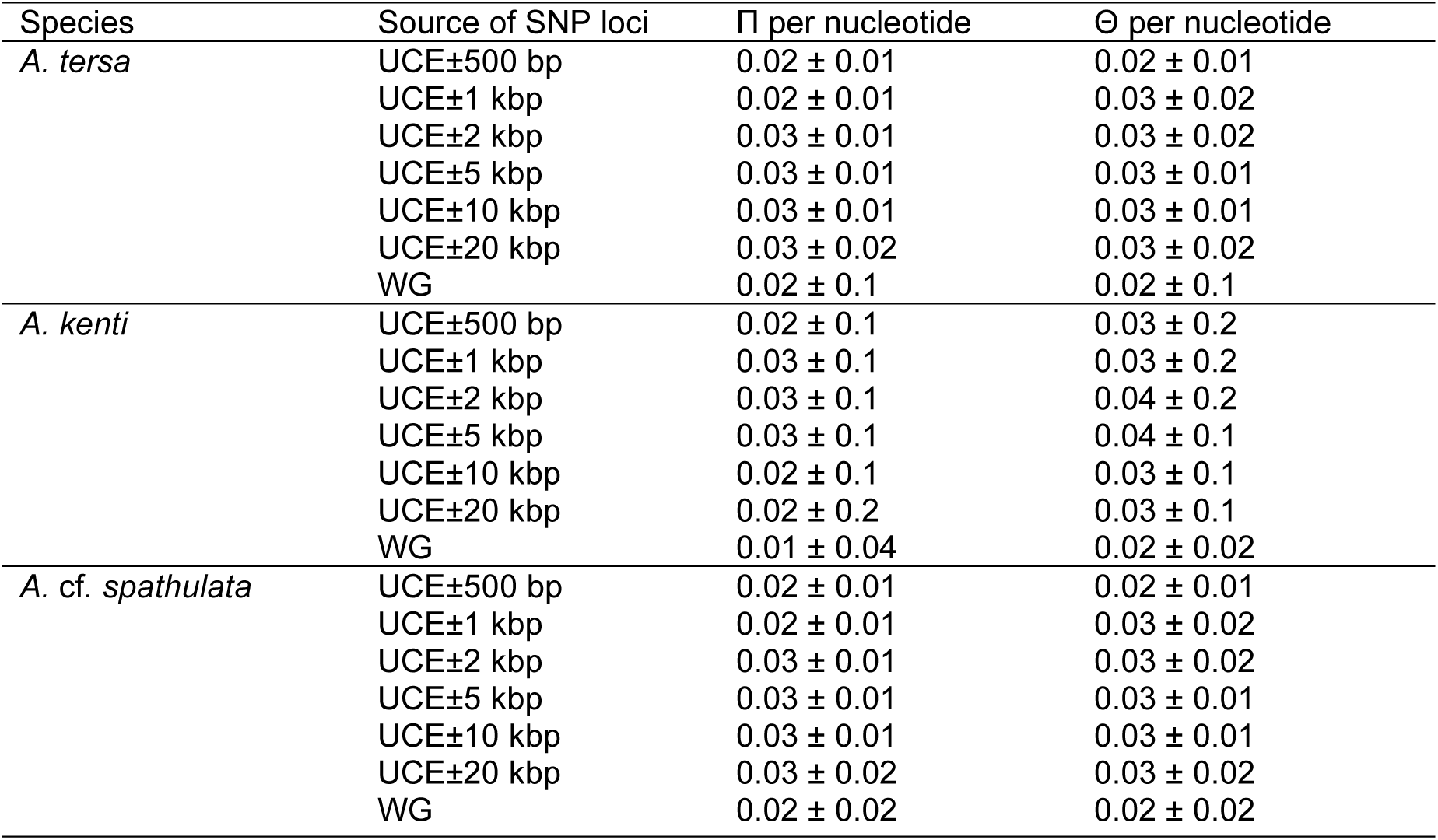
Measures of intraspecific variation calculated from SNP loci captured within extended UCE regions. SNPs derived from 200 UCE loci present on all taxa. Data are presented as means ± SD.

**ST4.** Measures of intraspecific variation calculated from SNP loci captured within extended UCE regions. SNPs derived from 100 UCE loci present on all taxa. Data are presented as means ± SD.

| Species | Source of SNP loci | $\Pi$ per nucleotide | $\Theta$ per nucleotide |
| --- | --- | --- | --- |
| *A. tersa* | UCE±500 bp | 0.02 ± 0.01 | 0.02 ± 0.01 |
| | UCE±1 kbp | 0.03 ± 0.01 | 0.03 ± 0.02 |
| | UCE±2 kbp | 0.03 ± 0.01 | 0.03 ± 0.02 |
| | UCE±5 kbp | 0.03 ± 0.01 | 0.03 ± 0.02 |
| | UCE±10 kbp | 0.03 ± 0.01 | 0.03 ± 0.01 |
| | UCE±20 kbp | 0.02 ± 0.02 | 0.03 ± 0.02 |
| | WG | 0.02 ± 0.1 | 0.02 ± 0.1 |
| *A. kenti* | UCE±500 bp | 0.02 ± 0.01 | 0.03 ± 0.02 |
| | UCE±1 kbp | 0.03 ± 0.01 | 0.03 ± 0.02 |
| | UCE±2 kbp | 0.03 ± 0.01 | 0.04 ± 0.02 |
| | UCE±5 kbp | 0.03 ± 0.01 | 0.04 ± 0.02 |
| | UCE±10 kbp | 0.02 ± 0.01 | 0.03 ± 0.01 |
| | UCE±20 kbp | 0.02 ± 0.02 | 0.03 ± 0.01 |
| | WG | 0.01 ± 0.04 | 0.02 ± 0.02 |
| *A. cf. spathulata* | UCE±500 bp | 0.02 ± 0.01 | 0.02 ± 0.01 |
| | UCE±1 kbp | 0.02 ± 0.01 | 0.03 ± 0.02 |
| | UCE±2 kbp | 0.03 ± 0.01 | 0.03 ± 0.02 |
| | UCE±5 kbp | 0.03 ± 0.01 | 0.03 ± 0.02 |
| | UCE±10 kbp | 0.03 ± 0.01 | 0.03 ± 0.01 |
| | UCE±20 kbp | 0.02 ± 0.02 | 0.03 ± 0.01 |
| | WG | 0.02 ± 0.02 | 0.02 ± 0.02 |

